# Accurate and scalable genome-wide ancestry estimation using haplotype clusters

**DOI:** 10.1101/2025.09.02.673718

**Authors:** Jonas Meisner

## Abstract

Unsupervised genome-wide ancestry estimation has been a staple in population and medical genetics for decades, and its importance continues to grow with the increasing number of large genetic cohorts of mixed ancestries. We propose an extension to the `hapla` framework that scales model-based ancestry estimation to unprecedented sample sizes by leveraging inferred haplotype clusters from phased genotype data. Our haplotype cluster-based approach is approximately 5× faster than the fastest model-free SNP-based approach on the harmonized Human Genome Diversity Project and 1000 Genomes Project dataset, while we further demonstrate it to be the most accurate method to date in an extensive simulation study. Our accurate ancestry estimates can help reduce health disparities and accelerate precision medicine efforts in the growing number of biobanks globally.

## Introduction

Ancestry estimation and population structure inference remain long-standing challenges in population and medical genetics. Genome-wide ancestry estimates in a population can capture evolutionary processes in its ancestral genetic sources [1], while also being instrumental in modern genome-wide association studies and polygenic score estimation to account for population stratification [2]. With the rapid growth of human biobank-scale genetic cohorts encompassing hundreds of thousands of samples [3, 4, 5], the need for accurate ancestry estimates has become increasingly urgent, as most modern human populations are or are becoming highly admixed. These large datasets introduce computational and methodological issues as well as bias concerns related to population structure, often leading to ad hoc simplifications such as predefined or self-reported ancestry categories. As a recent example, the All of Us research program [5] employs a supervised classification model, trained on data from the Human Genome Diversity Project [6] and the 1000 Genomes Project [7], to infer categorical ancestry labels from projected principal component scores. Additionally, admixed individuals are usually excluded from population-specific genome-wide association studies because standard methods cannot adequately handle or account for population structure and ancestry [8]. However, recent large-scale efforts show how tailored methods, which include admixed individuals, can increase statistical power [8, 9], thus demonstrating the importance of accurate ancestry estimates to ensure that valuable findings are not lost through the exclusion of genetic variation.

We recently developed the `fastmixture` [10] software to address a major gap in scaling accurate unsupervised model-based ancestry estimation to larger sample sizes. We demonstrated 30 × faster computational runtimes in comparison to the widely used `ADMIXTURE` [11] software, while maintaining equal or greater accuracy, and these runtime gains have further increased in recent versions of `fastmixture`. Another recent approach for scaling ancestry estimation is `SCOPE`, which utilizes alternating least-squares optimization to speed up inference [12], however, at the expense of accuracy [10].

Recent advances in efficiency and scalability of haplotype estimation algorithms [13, 14], including availability of massive reference panels [15], have enabled accurate phasing in large-scale genetic datasets. This has increased interest in leveraging haplotype information to, for example, infer local ancestry [16], detect fine-scale population structure [17], and even enhance statistical power in multi-ancestral genome-wide association studies [8]. We have also recently developed `hapla` [18], which performs efficient window-based haplotype clustering in phased genotype data. By generating local haplotype representations through inferred clusters, `hapla` aims to improve accuracy in heritability estimation and predictive performance in polygenic prediction.

Here we present a submodule of the `hapla` framework, denoted `hapla admix`, which integrates the fast mini-batch expectation-maximization algorithm in `fastmixture` with the haplotype cluster assignments in `hapla`. This enables accurate unsupervised model-based ancestry estimation for datasets at an unprecedented scale. Our approach facilitates the integration of accurate ancestry proportions into large cohorts, supporting multi-ancestral genome-wide association studies and polygenic score estimations without relying on predefined categorical population labels, thereby helping to reduce health disparities across ancestries.

## Methods

### Haplotype clustering

We generate haplotype cluster assignments using the `hapla cluster` submodule in `hapla` [18] from phased genotype data. Haplotypes are clustered in non-overlapping windows of fixed length, *B*, along a chromosome. Note that we use the term “haplotypes” to refer to the phased genotype data, with each sample having two phased copies. Within a window block of *B* SNPs, `hapla` assigns each haplotype to an inferred cluster representative based on Hamming distance. We note that in the special case of *B* = 1, `hapla` will simply generate the original phased genotype data, where each SNP is represented by two clusters corresponding to the major and minor allele.

Haplotype clusters are inferred in two steps: a discovery step generating *C*_0_ clusters, and a pruning and re-clustering step that iteratively removes haplotype clusters below a defined frequency threshold *δ*, yielding *C* ≤ *C*_0_ clusters. The resulting haplotype clustering produces a *W* × *N* × 2 haplotype cluster assignment tensor, **Z**, where *W* is the number of windows (across all chromosomes) and *N* is the number of samples. Each window has a unique number of inferred clusters, such that *z*_*wih*_ ∈ {1, …, *C*^(*w*)^} is the haplotype cluster assignment in a given window *w* for sample *i* and haplotype *h*, and *C*^(*w*)^ is the number of haplotype clusters in window *w*. We will investigate the effect of *B* and *δ* on the accuracy and computational runtime in unsupervised ancestry estimation using simulated datasets, as different window sizes and cluster frequency thresholds may capture different aspects in terms of ancestry tracts and genomic diversity, respectively.

### Ancestry estimation

#### Likelihood model

We utilize the haplotype cluster assignment matrix from the previous step to perform model-based ancestry estimation. Our approach, denoted `hapla admix`, can be viewed as an extension of the original `ADMIXTURE` [11] likelihood model to account for multi-allelic SNPs in haploid data, where SNPs and alleles in the original model correspond to windows and haplotype clusters in our context, respectively, which has also previously been derived in `HaploNet` [19]. The likelihood model is defined as follows for a predefined number of ancestral components *K*, assuming independence across windows, samples, and haplotypes:

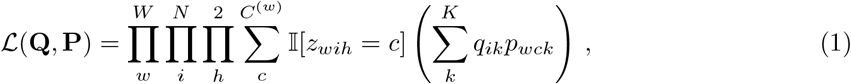

where 𝕀[·] is the indicator function, *q*_*ik*_ is the *k*-th ancestry proportion of sample *i*, and *p*_*wck*_ is the ancestral haplotype cluster frequency in the *k*-th ancestry component of cluster *c* in window *w*. The sample ancestry proportions, **Q** ∈ [0, 1]^*N*×*K*^, are constrained such that 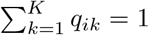 for all *i*, and the ancestral haplotype cluster frequencies, **P** ∈ [0, 1]^*W* ×*C*×*K*^, are similarly constrained such that 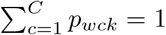 for all *w* and *k*.

### Maximum likelihood and accelerated optimization

We estimate the ancestry proportions and ancestral haplotype cluster frequencies using a maximum likelihood approach with an expectation-maximization (EM) algorithm. Because the EM algorithm notoriously suffers from slow convergence in high-dimensional data [20], we adapt the acceleration approach from `fastmixture`, which combines a quasi-Newton acceleration scheme with mini-batch updates. The update rules for **Q** and **P** at iteration *t* are defined below, representing a slight adaptation of the EM updates derived in `HaploNet` for its haplotype cluster likelihoods adjusted to our haplotype cluster assignments:

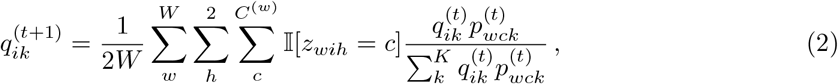

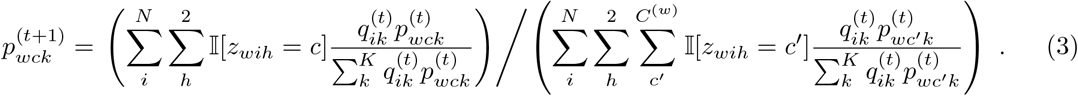

The mini-batch EM updates are performed across windows in our implementation. To further accelerate the convergence of the EM algorithm, we adopt the singular value decomposition (SVD) initialization procedure from `fastmixture`. Specifically, we utilize a randomized SVD algorithm [21] to generate a latent subspace in which we employ an alternating least-squares (ALS) procedure, following `ALStructure` [22] and `SCOPE` [12], to obtain better starting points for **Q** and **P** [10]. A description of the SVD initialization procedure is detailed in the Supplementary Material.

We terminate the iterative updates of the EM algorithm when the difference in scaled loglikelihood units between successive iterations falls below a threshold *ϵ* = 1.0 × 10^−9^, such that convergence is determined as:

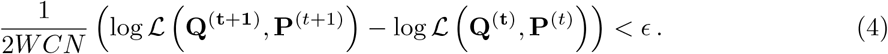

### Implementation

The `hapla` framework (v0.32.2), including `hapla admix`, is implemented as a Python command-line tool with a Cython backend. Haplotype cluster assignments from the `hapla cluster` submodule are stored in a binary file format (unsigned 8-bit integers) for efficient and fast integration with the `hapla admix` submodule. During the SVD initialization, we only expand the haplotype cluster assignments in user-defined sized chunks, as we perform a batched version of randomized SVD to reduce the memory footprint with only a minor runtime penalty. The ancestry estimation algorithm never expands the haplotype cluster assignment matrix, such that the memory footprint of `hapla admix` is mostly dominated by the input matrix of *W* × *N* × 2 bytes, where *W* is the number of windows, *N* is the number of samples, and each sample has two haplotypes.

We have further implemented projection and supervised modes in `hapla admix` to enhance its usability. The projection mode allows for rapid haplotype clustering and ancestry estimation in a target dataset based on the inferred haplotype clusters and their ancestral cluster frequencies from a reference dataset. These options facilitate the integration of ancestry estimates in workflows, such as polygenic score estimation, where external validation is standard.

### Simulated data

We simulate human-based genetic datasets using the population genetics frameworks `msprime` [23] and `tskit` [24]. We construct two demographic models of differing complexity for ancestry simulations, where we simulate genomic segments of 200 megabases assuming an uniform recombination rate of 1.28 × 10^−8^ per base pair per generation [25] and a mutation rate of 1.44 × 10^−8^ per base pair per generation [26] in both.

The first demographic model (Scenario 1) is based on the American admixture model [27], which emulates an out-of-Africa event with African, Asian, and European source populations (*K* = 3), including an admixed population from a recent admixture event 12 generations ago. The second demographic model (Scenario 2) includes five source populations (*K* = 5) and three admixed populations arising from both recent and deeper admixture events (10, 20, and 30 generations ago). For both scenarios, we simulate three datasets of increasing sample sizes (*N* = {2,000, 8,000, 32,000}), yielding a total of six datasets to evaluate the performance and scalability of different ancestry estimation methods in moderate to large-scale datasets of varying complexity. We denote the different sample sizes A, B, and C for 2,000, 8,000, and 32,000, respectively, such that Scenario 1A refers to Scenario 1 with 2,000 samples. We perform SNP filtering with a minor allele frequency threshold of 0.05 in all datasets.

We use the `tspop` [28] Python package to track ancestry tracts in the tree sequences from the `msprime` simulations. We insert a census event 100 generations back in time to obtain ground truth ancestry proportions. Therefore, gene flow between source populations is not allowed until after the census event has occurred to avoid assigning the wrong ancestral sources to the simulated ancestry tracts because of prior gene flow. The demographic models are visualized in Supplementary Figures 1 and 2, and general details about the six generated datasets are provided in the Supplementary Material.

### Harmonized HGDP and 1KGP data resource

We make use of the high-quality harmonized phased genotype dataset [29], which combines the Human Genome Diversity Project [6] and the 1000 Genomes Project [7]. It contains 4,094 whole genomes from 80 populations worldwide and is the largest publicly available human genetic dataset, making it a common reference panel for phasing and imputation. We remove related samples based on a kinship coefficient threshold of 0.05, while optimizing for the total sample size, using the KING-robust estimator [30] implemented in `PLINK` (v2.0.0a) [31], leaving 3,406 unrelated samples. After applying a standard minor allele frequency threshold of 0.05, a total of 5,870,342 diallelic SNPs are retained.

## Results

We evaluate our haplotype cluster-based approach, `hapla admix`, against two scalable SNP-based methods for unsupervised genome-wide ancestry estimation. `SCOPE` [12] uses a model-free alternating least-square (ALS) approach to scale ancestry estimation to biobank-sized datasets, however, as previously shown, its scalability comes at the cost of accuracy in larger and complex scenarios [10]. We also compare our haplotype cluster-based approach to its SNP-based alternative, `fastmixture` [10] (v1.1.1), to directly assess the gains in scalability and accuracy from leveraging haplotype information. For simplicity, we refer to `hapla admix` as `hapla`. Computational runtimes of all software are measured across all scenarios on an Intel(R) Xeon(R) Gold 6152 CPU @ 2.10GHz, utilizing 44 threads.

As the input data and its format differ between `hapla` and the SNP-based methods, direct comparison of estimated ancestry proportions is only possible in the simulated datasets, where we report the root mean square error (RMSE) against the ground truth. For the empirical dataset, we report the log-likelihoods across multiple runs for all three methods, as these reflect the overall variability of their solutions. Descriptions of the evaluation metrics are provided in the Supplementary Material.

### Ancestry estimation in simulation study

We have simulated six datasets from two demographic models of differing complexity, each at three different sample sizes of 2,000, 8,000, and 32,000, denoted A, B, and C, respectively. The first scenario is a custom adaptation of the American admixture model [27], which emulates an out-of-Africa event with three source populations (*K* = 3), including an admixed population. The second demographic model includes two bottleneck events and consists of five source populations (*K* = 5) and three admixed populations.

### Effect of window size and cluster frequencies

We investigate the effect of window size, *B*, and cluster frequency threshold, *δ*, in the haplotype clustering step of `hapla` [18] to identify an optimal set of parameters that minimizes the RMSE to the ground truth ancestry proportions in the ancestry estimation step in the simulated datasets. We also report the computational runtimes of the different combinations for both the haplotype clustering step and the ancestry estimation step. Ancestry estimation is evaluated for *B* = {8, 16, 32} and *δ* = {0.005, 0.01, 0.05}.

We immediately note that the lowest cluster frequency threshold, *δ* = 0.005, produces the most accurate ancestry estimates for all window sizes in all simulated datasets (Supplementary Table 1). This threshold also yields the fastest computational runtimes in the clustering step, as `hapla` has to perform fewer iterations of the cluster pruning and re-clustering procedure, while adding only a minimal runtime penalty of 0.04 minutes on average for retaining the extra clusters in the ancestry estimation step compared to *δ* = 0.05 (Supplementary Figure 6). Ancestry plots for Scenario 1A and 2A with the three window sizes using *δ* = 0.005 are shown in Supplementary Figures 3 and 4, respectively.

Increasing the window size results in substantial reductions in computational runtimes for ancestry estimation, as the computational complexity of our approach scales linearly with the number of windows. For *δ* = 0.005, the runtime decreases by approximately 65% on average between *B* = 8 and *B* = 32 (Supplementary Table 5). In terms of accuracy, *B* = 8 performs worst in all scenarios, indicating that larger window sizes better model the ancestry tracts in the simulated datasets (Supplementary Table 1). Comparing window sizes, *B* = 16 and *B* = 32, in Scenario 1, *B* = 16 is slightly more accurate for 1A and 1B, with RMSE values of 0.00473 and 0.00453 versus 0.00490 and 0.00459 for *B* = 32. However, for the larger scenario (1C), *B* = 32 has a lower RMSE of 0.00464 compared to 0.00469 for *B* = 16. In the more complex simulation Scenario 2 with five source populations, *B* = 32 outperforms the other two window sizes for *δ* = 0.005 with RMSE values of 0.00354, 0.00343, and 0.00340 for 2A, 2B, and 2C, respectively, whereas *B* = 16 has values of 0.00358, 0.00349, and 0.00349. Overall, *B* = 16 and *B* = 32 perform nearly equally across all scenarios for *δ* = 0.005, while *B* = 32 offers substantially lower computational runtimes.

Based on these results, we select *B* = 32 and *δ* = 0.005 as the optimal combination for ancestry estimation using haplotype clusters in our human-based simulation study, balancing both accuracy and computational runtime. This parameter set is used in comparisons with the SNP-based approaches and in the empirical data application.

### Comparison to other approaches

We evaluate the performance of `hapla` against the SNP-based approaches, `fastmixture` and `SCOPE`, in the simulations. Overall, our haplotype cluster-based approach is significantly more accurate than the two SNP-based approaches across all datasets, while `fastmixture` consistently outperforms `SCOPE` when comparing only the two SNP-based approaches (Table 1). `hapla` shows reductions in RMSE of approximately 71% compared to `SCOPE` and 32% relative to `fastmixture`. `SCOPE` introduces uniform noise in its ancestry estimates, as previously reported [10], which is clearly visible in the unadmixed populations in Figure 1 and Supplementary Figure 5 for Scenario 1A and Scenario 2A, respectively. The noise is less visually pronounced in admixed populations, where it may be masked since most of the ancestry components are represented.

**Table 1:** Root mean square error (RMSE) between the ground truth and estimated ancestry proportions in the simulated data. Lower values indicate better performance. Reported values are the mean RMSE value across five runs, and the standard deviations are omitted as all are *<* 1.0 × 10^−5^. The best-performing method in each scenario is highlighted in bold. `hapla` results use *B* = 32 and *δ* = 0.005. The source data for the table can be found in the Supplemental Data file.

| Method | Scenario 1 |  |  | Scenario 2 |  |  |
| --- | --- | --- | --- | --- | --- | --- |
|  | A | B | C | A | B | C |
| SCOPE | 0.01512 | 0.01525 | 0.01486 | 0.01243 | 0.01272 | 0.01260 |
| fastmixture | 0.00651 | 0.00647 | 0.00646 | 0.00532 | 0.00540 | 0.00546 |
| hapla | **0.00490** | **0.00459** | **0.00464** | **0.00354** | **0.00343** | **0.00340** |

**Figure 1:**
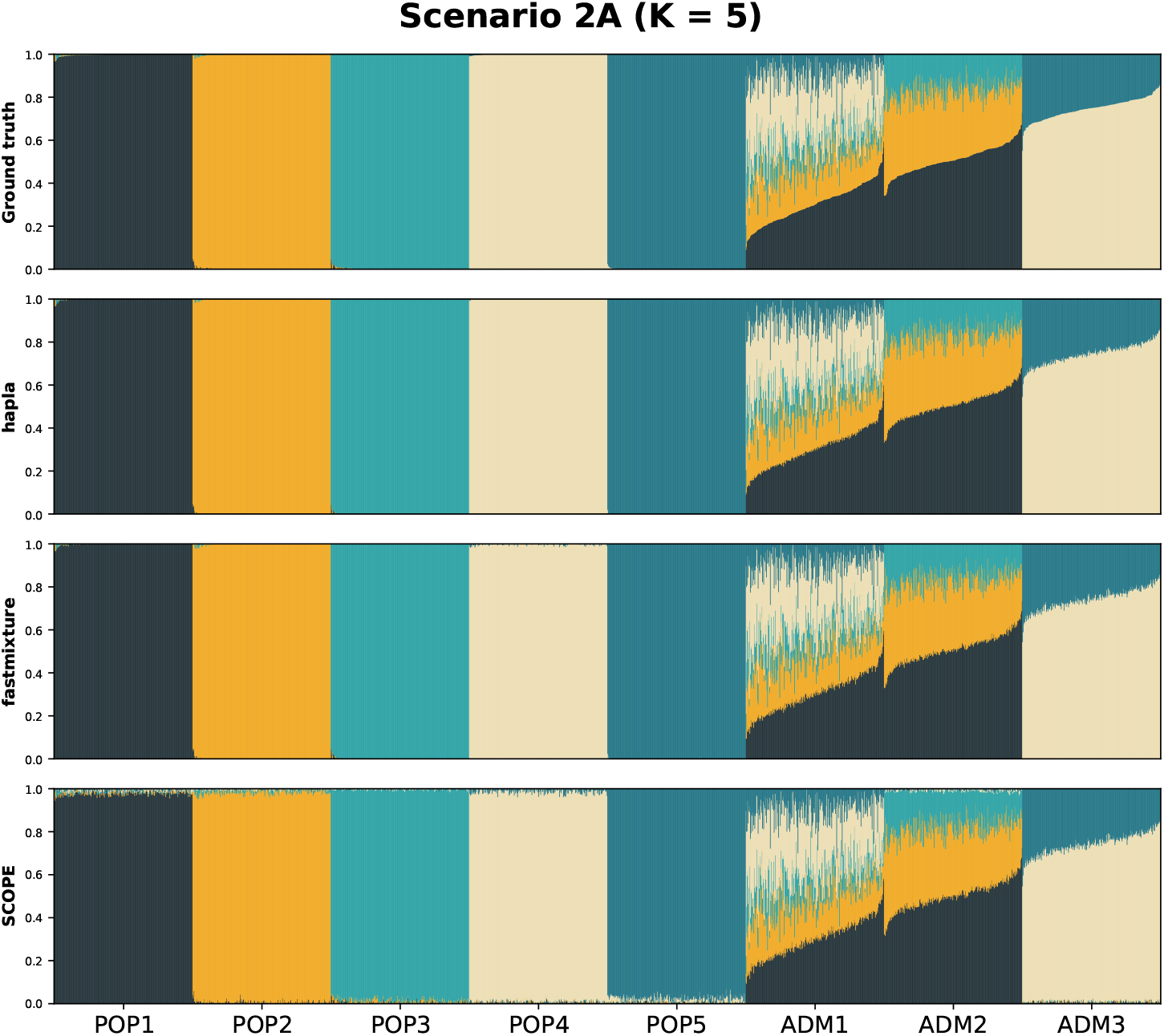
Ancestry plots of the simulated Scenario 2A of 2,000 samples for *K* = 5. The top ancestry plot displays the ground truth ancestry proportions, obtained from simulated ancestry tracts, followed by the estimates of the three evaluated methods.

We also compare the RMSE values using only the admixed populations to investigate potential differences in accuracy. The same trend observed for all populations holds, where `hapla` is more accurate than the other two methods (Supplementary Table 2). In this comparison, `hapla` reduces RMSE values by approximately 49% compared to `SCOPE` and 36% relative to `fastmixture`. The model-based approaches estimate ancestry proportions of the unadmixed populations slightly better than those of the admixed populations, whereas `SCOPE` shows a negligible difference between them.

Finally, we compare the computational runtimes across methods in the simulated datasets. `hapla` not only outperforms the SNP-based approaches in accuracy, but also achieves significantly faster runtimes (Figure 3). In Scenario 1 for *K* = 3, `hapla` is 1.6× and 12.3× faster than `SCOPE` and `fastmixture`, respectively, averaged across the three sample sizes, and in Scenario 2 for *K* = 5, it is 2.4× and 12.5× faster.

### Application to the harmonized HGDP and 1KGP data

We next apply our method to the harmonized Human Genome Diversity Project (HGDP) and 1000 Genome Project (1KGP) dataset [29]. After standard filtering, the dataset consists of 3,406 individuals and 5,870,342 diallelic SNPs. Ancestry is estimated across a range of ancestry components (*K* ={5, 6, 7}). The individuals represent populations from around the globe, where the following seven metadata labels have been defined in the dataset: Africa (AFR), America (AMR), Central South Asia (CSA), East Asia (EAS), Europe (EUR), Middle East (MID), and Oceania (OCE).

We visualize the ancestry plots for the three different evaluated methods using *K* = 5, *K* = 6, and *K* = 7 in Supplementary Figures 7 and 8, and Figure 2, respectively. Overall, `SCOPE` exhibits a pronounced layer of uniform noise in its ancestry estimates compared to the simulated datasets, and this noise increases with higher numbers of ancestry components. For all three values of *K*, `SCOPE` fails to capture any unadmixed individuals, except for a few individuals in the American component. `fastmixture` generates similar ancestry proportions to `hapla` for *K* = 5, however, the two methods start to differ slightly for *K* = 6 and *K* = 7, where `hapla` captures a more distinct ancestry component for the OCE individuals, which appears to also be captured in the CSA individuals for the SNP-based approaches.

**Figure 2:**
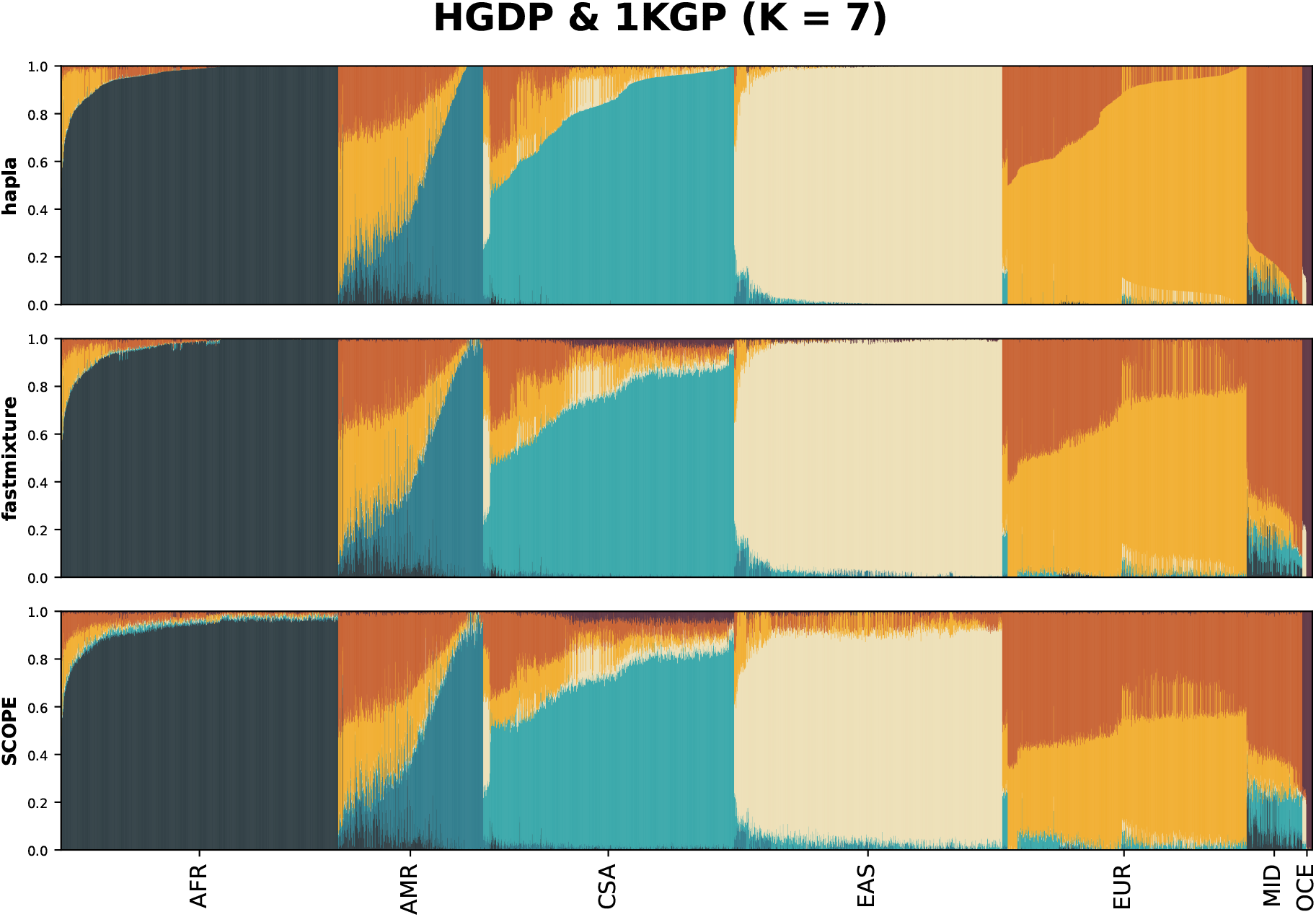
Ancestry plots of 3,406 individuals in the harmonized HGDP and 1KGP dataset for *K* = 7 using the three evaluated methods. The order of the individuals is based on metadata labels and the ancestry estimates of `hapla`.

A limitation observed in both SNP-based approaches is the absence of individuals with near 100% ancestry in the CSA ancestry component across all scenarios, and in the MID ancestry component for the *K* = 7 scenario. This is a necessary condition to have an identifiable admixture model, as there needs to exist an “anchor point” for each ancestry component [22, 32]. While the overall ancestry structure captured by `hapla` is reflected in the SNP-based methods, these approaches lack the resolution to fully model the Central South Asian and Middle Eastern components, which are both directly modeled by the haplotype cluster-based approach. Due to the noise layer in the ancestry estimates of `SCOPE`, it rarely satisfies the anchor condition in any of the three *K* scenarios as mentioned above, including the scenarios of the simulated study.

The log-likelihoods for each method are reported in Supplementary Table 3, showing little variation across the five runs, which indicates that all methods find stable solutions. The variation is larger for `SCOPE`, but still negligible and likely a result of it not maximizing the likelihood model but a least-squares objective instead. From a model selection viewpoint, we report the Akaike information criterion (AIC) [33] based on the log-likelihoods in Supplementary Table 4 to assess the optimal number of ancestry components. AIC values indicate *K* = 7 as the most appropriate model across all methods, whereas the original study [29] identifies *K* = 6 as the most appropriate model based on a cross-validation metric.

We also evaluate the computational runtimes for ancestry estimation across various numbers of ancestry components. Consistent with the simulation study, `hapla` showcases much faster runtimes than the two SNP-based approaches, averaging 5.3 × faster than `SCOPE` and 19.7 × faster than `fastmixture` across the three evaluated values of *K* (Figure 3). Focusing on the haplotype clustering step in `hapla`, it only takes approximately 2 minutes to process the largest chromosome in the dataset (chr2) with 476,184 SNPs (Supplementary Figure 6), and the runtime can be amortized across all chromosomes by parallel processing.

**Figure 3:**
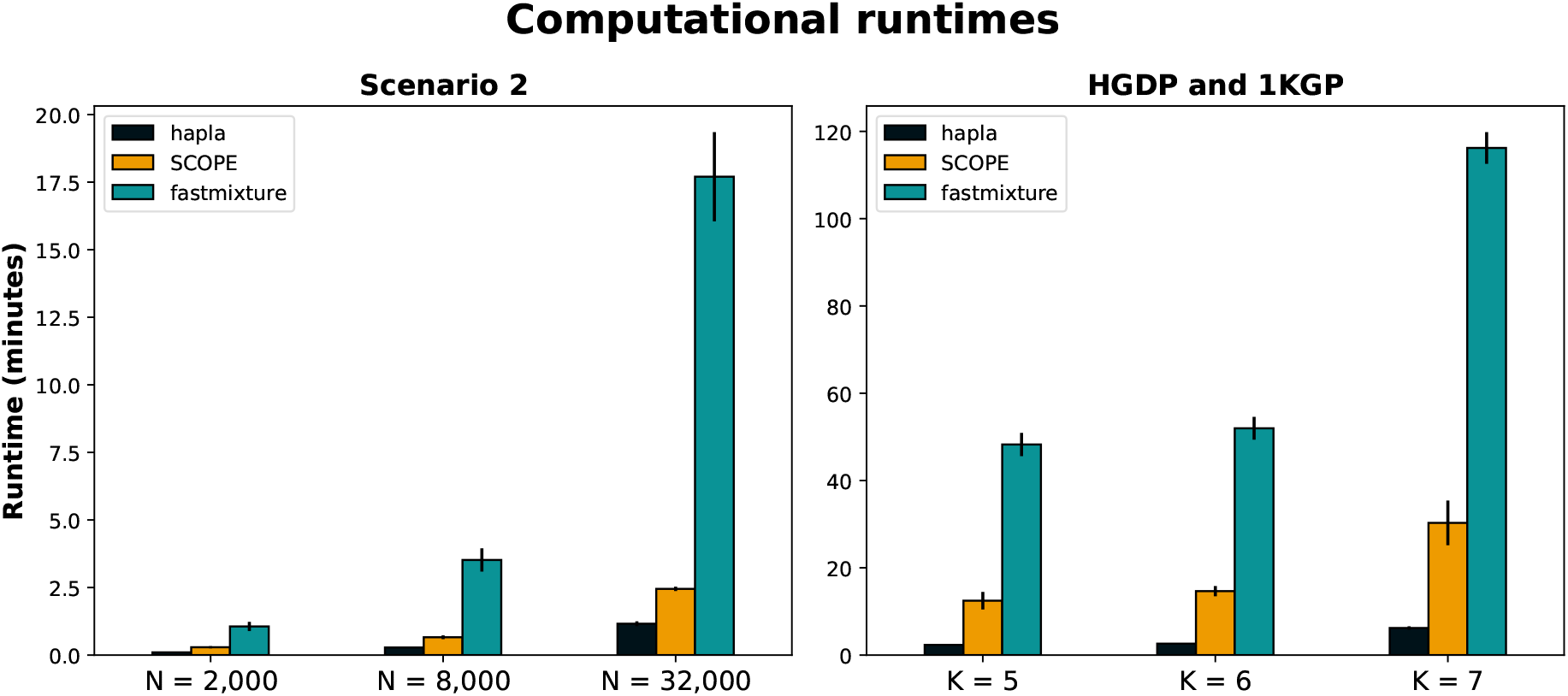
The computational runtimes *in minutes* for performing unsupervised ancestry estimation using 44 threads. Left plot: the second simulation scenario (*K* = 5) for various sample sizes. Right plot: the harmonized HGDP and 1KGP dataset consisting of 3,406 individuals and 5,870,342 SNPs, for various numbers of ancestry components. The means and standard deviations (error bars) are shown across five runs. Lower values indicate faster runtimes. The source data for the figure can be found in the Supplemental Data file.

## Discussion

We have proposed a model-based approach for unsupervised genome-wide ancestry estimation using haplotype clusters, `hapla admix`, implemented within the `hapla` framework. The method leverages haplotype cluster assignments from a window-based clustering procedure, enabling indirect use of the haplotype information from phased genotype data in ancestry estimation. We adopt a likelihood model where we compute maximum likelihood ancestry estimates through a mini-batch accelerated expectation maximization (EM) algorithm.

We demonstrate that `hapla` is both the most accurate and the fastest method for unsupervised ancestry estimation in simulations across multiple sample sizes and two scenarios of differing complexity. Both in simulations and in the harmonized Human Genome Diversity Project (HGDP) and 1000 Genomes Project (1KGP) dataset, we provide further evidence that the currently fastest model-free approach, `SCOPE`, based on alternating least-squares optimization, is inferior in accuracy compared to maximum likelihood approaches. Across all analyses in our study, `SCOPE` introduces a noise layer that increases in magnitude for larger and more complex ancestry scenarios. Despite its lower accuracy, the speed of the alternating least-squares optimization enables scaling to biobank-sized datasets. In our method, we therefore still utilize a model-free approach to generate noisy ancestry estimates as starting points, which accelerates the convergence of our maximum likelihood estimation, as previously reported in the `fastmixture` study. The SNP-based variant of our proposed method, `fastmixture`, also scales model-based ancestry estimation to large datasets, but it will not be feasible to run the method at biobank-scale, and it lacks the resolution gained in the haplotype cluster-based approach, as shown in the empirical data application. By combining the inferred haplotype clusters and the mini-batch EM algorithm of `fastmixture`, `hapla` scales model-based ancestry estimation to unprecedented sample sizes, while maintaining high accuracy and capturing fine-grained ancestry signals.

A limitation of our approach is the need for phased genotype data, which may not be readily available or highly accurate for most genetic datasets. The utilization of haplotype information introduces additional pipeline steps regarding haplotype clustering, including decisions on window size and cluster frequency threshold, which may be dataset or organism-specific. In this study, we explored the basis of unsupervised ancestry estimation in human sequencing data, and the parameter decisions may vary with differing SNP densities or mutation rates. Nevertheless, *all* tested parameter combinations in `hapla` outperform the SNP-based approaches across all simulated datasets, demonstrating the robustness of our haplotype cluster-based approach. Therefore, our results further highlight the potential and opportunities that haplotype information can provide to enhance a range of analyses in population and medical genetics.

## Code and data availability statement

The `hapla` framework, including its haplotype clustering and ancestry estimation submodules, is freely available on GitHub (https://github.com/Rosemeis/hapla). All analyses are based on open and public datasets. The processed datasets, results, and scripts to reproduce the data generation process are available on Zenodo (https://doi.org/10.5281/zenodo.17026644). The harmonized phased genotype dataset of the Human Diversity Genome Project and the 1000 Genomes Project is publicly available at https://gnomad.broadinstitute.org/downloads#v3-hgdp-1kg.

## Acknowledgments

J.M. is supported by the Novo Nordisk Foundation (NNF24SA0093895). We thank Alba Refoyo Martinez for fruitful discussions and comments.

## Author contributions

J.M. conceived the study, derived the method, and implemented the software. J.M. performed all analyses and wrote the manuscript.

## Supplementary Material

### SVD initialization

We adapt the singular value decomposition (SVD) and alternating least-squares (ALS) initialization approach from `fastmixture` [10] to the haplotype cluster assignment setting in `hapla admix`, providing improved starting points for **Q** and **P**. For simplicity of notation, we assume that each window has the same number of inferred haplotype clusters *C*. To apply SVD, we first transform the haplotype cluster assignment tensor, **Z** ∈ {1, …, *C*} ^*W* ×*N*×2^ into a two-dimensional matrix. Specifically, we count the cluster assignments per sample for each of the inferred haplotype clusters, **Z**_*A*_ ∈ {0, 1, 2} ^*L*×*N*^, as also performed in the original `hapla` study for constructing a genome-wide relationship matrix [18]. Here, *L* = *WC* denotes the total number of haplotype clusters. We then perform SVD on the centered *Z*_*A*_ matrix to extract the top *D* = *K* − 1 singular values and singular vectors, 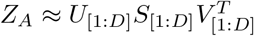.

The individual haplotype cluster frequency of sample *i* at haplotype cluster *l* can be approximated via SVD as 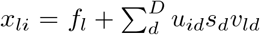, which we use in an ALS approach, as in `ALStructure` [22] and `SCOPE` [12], to obtain **Q** and **P**. The iterative ALS approach optimizes the following least-squares objective:

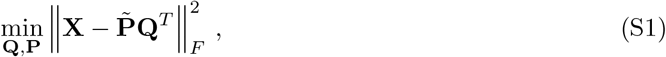

where 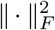 denotes the Frobenius norm, **X** is the individual haplotype cluster frequency matrix, and 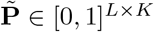 is a reshaped representation of the ancestral haplotype cluster frequency matrix, **P**, to make computations feasible. After each update step of **Q** and **P** in the ALS approach, we normalize them based on their individual constraints. The iterative ALS approach is terminated when the root mean square error between two successive iterations of **Q** is below 1.0 × 10^−4^.

### Evaluation metrics

#### Root mean square error

The root mean square error (RMSE) between the ground-truth ancestry proportions **Q**^*^ and the estimated ancestry proportions 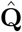 is defined as:

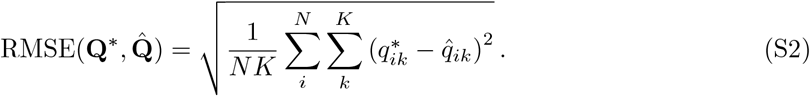

To avoid performance differences between methods due to numerical optimization, we clamp the estimated ancestry proportions between 0.00001 and 0.99999 and re-normalize the proportions to sum to one, such that comparisons are performed on equal scales.

#### Log-likelihood

In our haplotype cluster–based approach, the log-likelihood is defined as:

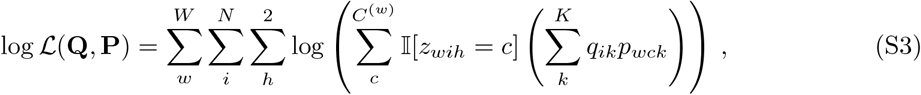

and in the SNP-based approaches:

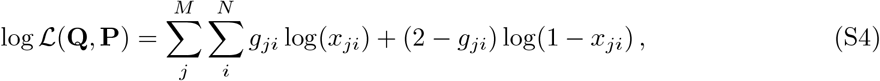

where *M* is the number of SNPs, *g*_*ji*_ is the diallelic genotype, and 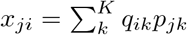 represents the individual allele frequency for SNP *j* in sample *i*.

#### Information criterion

The Akaike information criterion (AIC) [33] is widely used for model selection and is defined as 2*θ*− 2 log *L*(**Q, P**), wh e re *θ* is the t otal nu mber of parameters. In our haplotype cluster-based approach,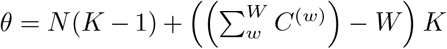, while in the SNP-based approaches *θ* = *N* (*K* − 1) + *MK*.

### Simulation study

Datasets are simulated using `msprime` [23]. Demographic models and scripts for reproducing the simulation scenarios, including the phased genotype datasets, are available on Zenodo (https://doi.org/10.5281/zenodo.17026644).

`msprime` **options**

- Segment length: 2.0 × 10^8^ bases
- Recombination rate: 1.28 × 10^−8^ per base pair per generation [25]
- Mutation rate: 1.44 × 10^−8^ per base pair per generation [26]

#### Scenario 1

- 3 source populations (AFR, EUR, ASN)
- 1 admixed population 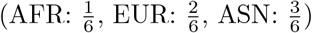

### Number of samples and SNPs (minor allele frequency threshold of 0.05)

**Number of samples and SNPs (minor allele frequency threshold of 0.05):**
|  | Scenario 1 |  |  |
| --- | --- | --- | --- |
|  | A | B | C |
| $N$ | 2,000 | 8,000 | 32,000 |
| $M$ | 244,356 | 241,652 | 241,226 |

#### Scenario 2

- 5 source populations (POP1, POP2, POP3, POP4, POP5)
- 1 admixed population 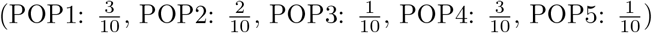
- 1 admixed population 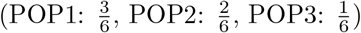
- 1 admixed population 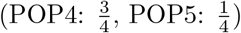

### Number of samples and SNPs (minor allele frequency threshold of 0.05)

**Number of samples and SNPs (minor allele frequency threshold of 0.05):**
|  | Scenario 2 |  |  |
| --- | --- | --- | --- |
|  | A | B | C |
| $N$ | 2,000 | 8,000 | 32,000 |
| $M$ | 284,616 | 281,330 | 281,090 |

## Supplementary Figures

**Supplementary Figure 1:**
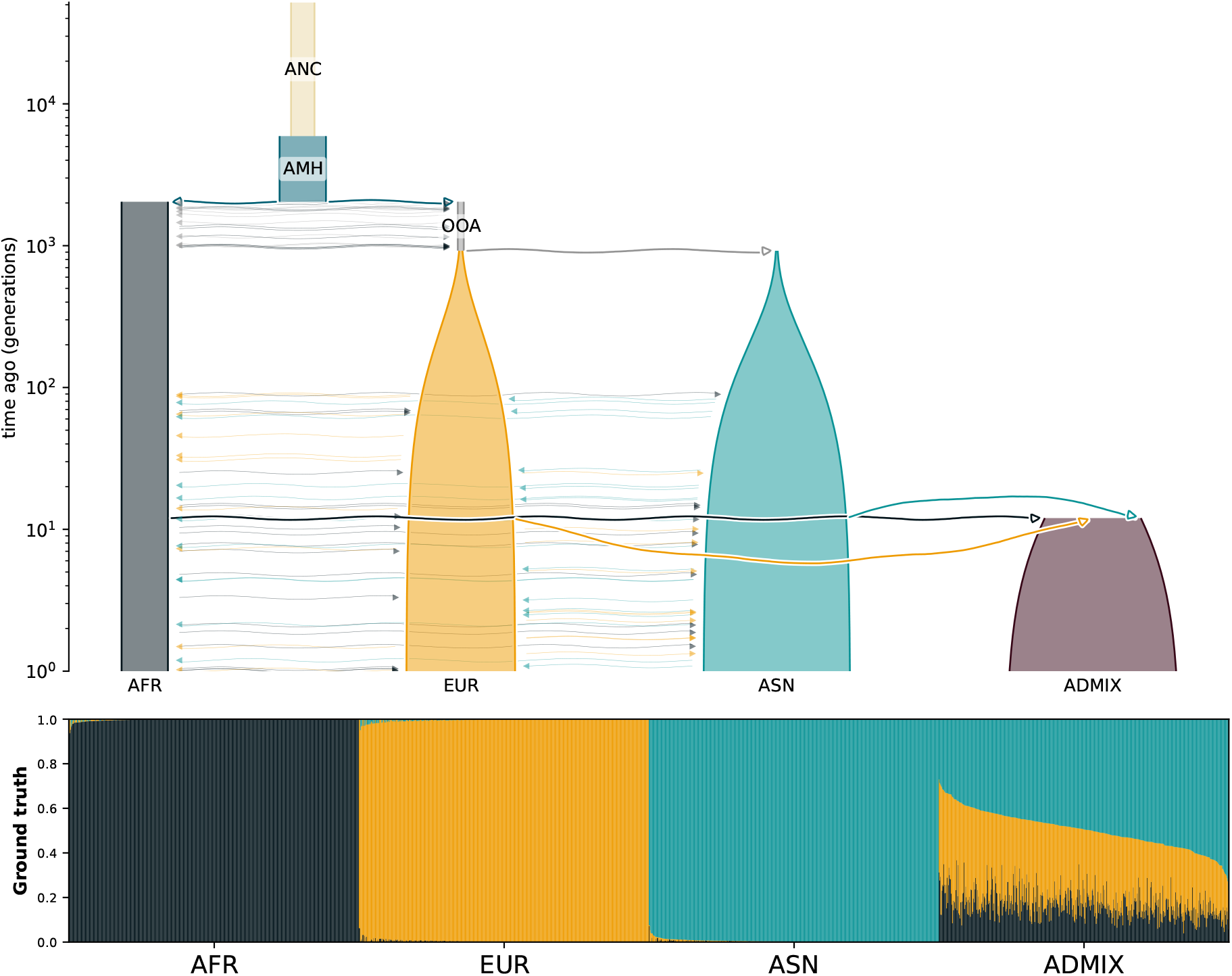
Demographic model of *Scenario 1* which is based on the American admixture model [27] with three source populations (*K* = 3) and one admixed population in the top plot. Note that we have disallowed gene flow between the source populations (AFR, ASN, EUR) until the census event has occurred (100 generations ago). Visualized using `demesdraw` [34]. The bottom plot shows the ground truth ancestry proportions in the simulated dataset of 2,000 samples obtained from the traced ancestry tracts using `tspop` [28].

**Supplementary Figure 2:**
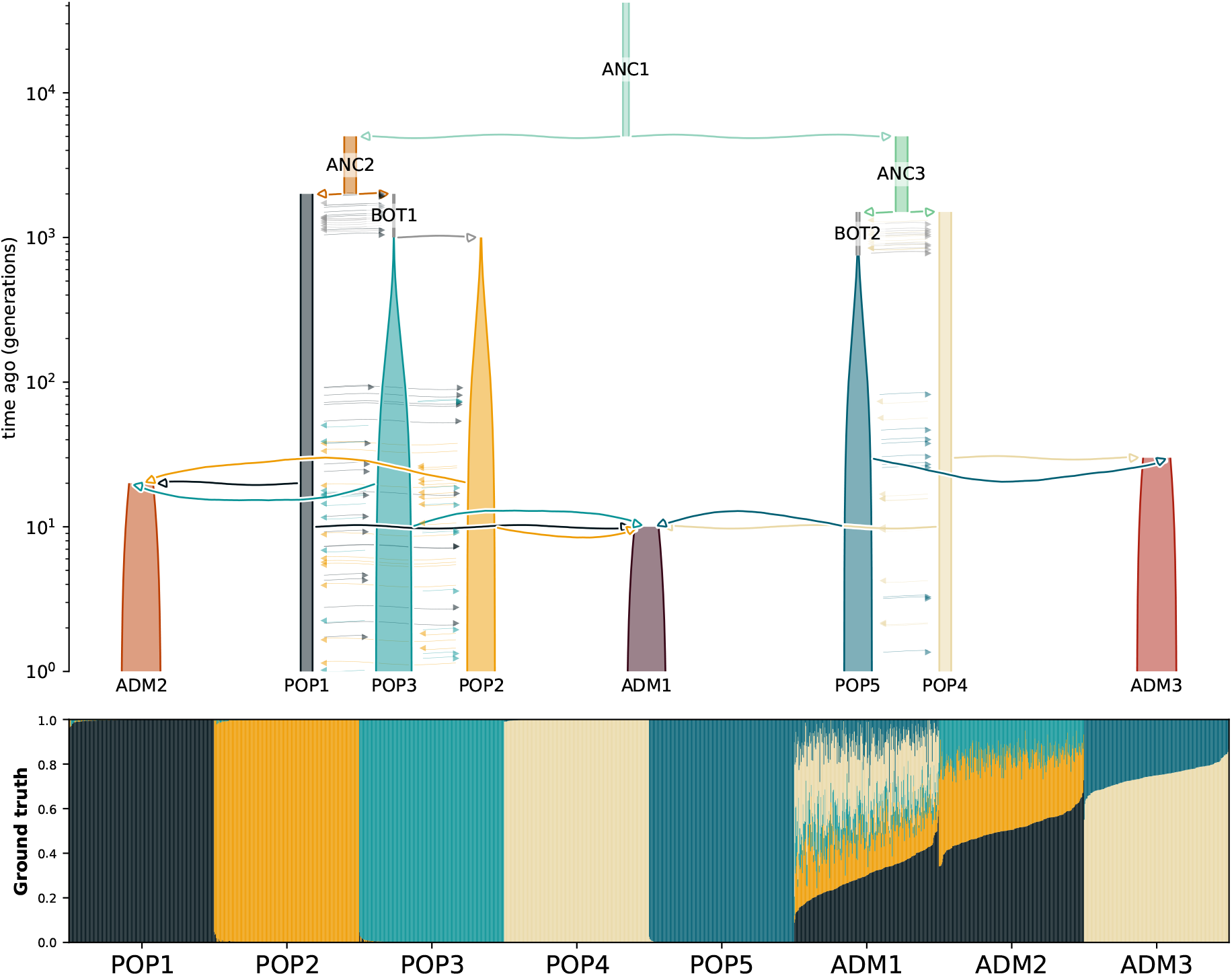
Demographic model of *Scenario 2*, based on a custom model with five source populations (*K* = 5) and three admixed populations (top plot). Gene flow was disallowed between the source populations (POP1, POP2, POP3, POP4, POP5) until the census event had occurred (100 generations ago). Visualized using `demesdraw` [34]. The bottom plot shows the ground truth ancestry proportions for 2,000 simulated samples, obtained from the traced ancestry tracts using `tspop` [28].

**Supplementary Figure 3:**
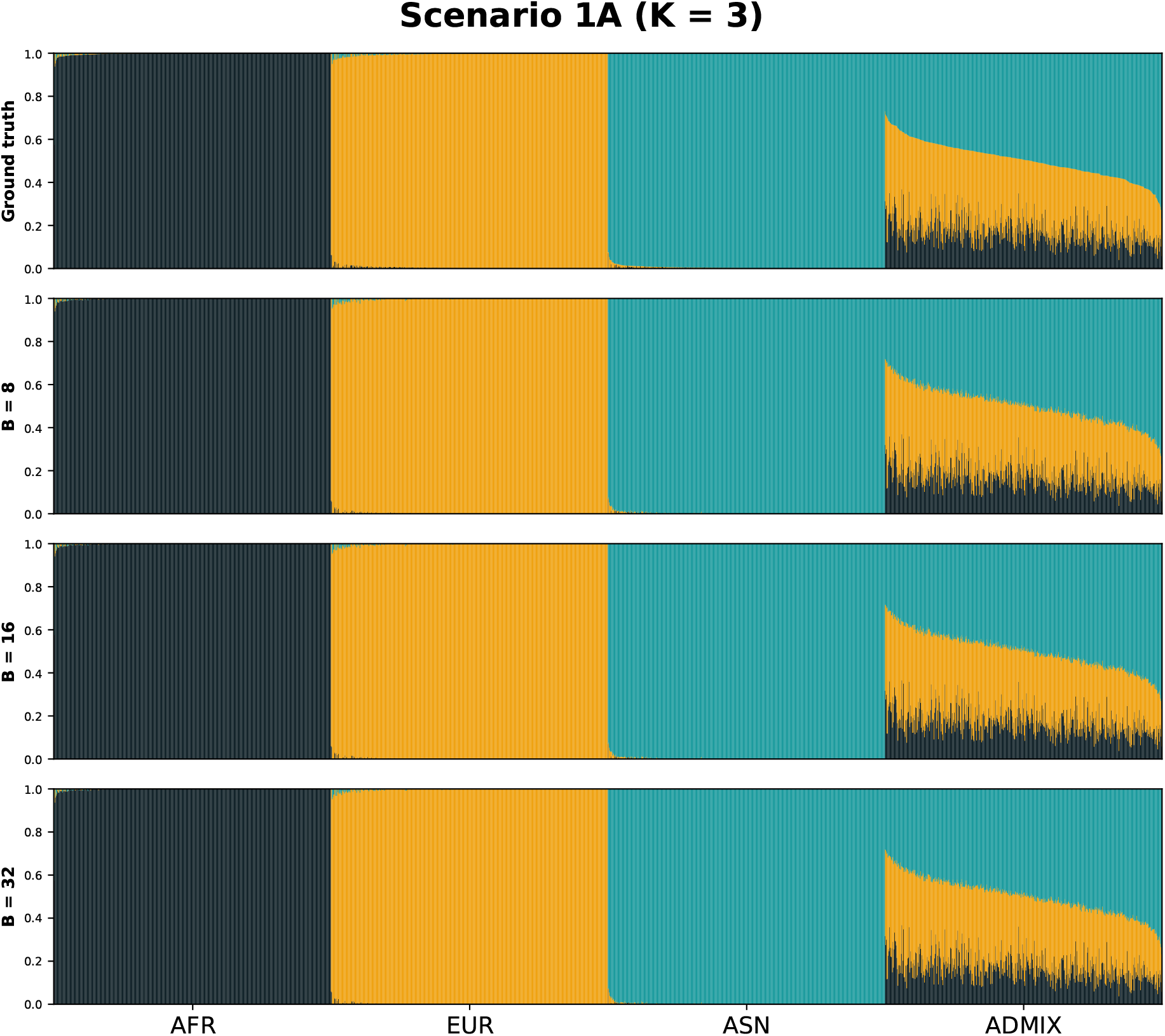
Ancestry plots for the simulated *Scenario 1A* of 2,000 samples for *K* = 5. The top ancestry plot displays the ground truth ancestry proportions from simulated ancestry tracts, followed by the estimates of three different window sizes in `hapla`, *B* = {8, 16, 32}, with *δ* = 0.005.

**Supplementary Figure 4:**
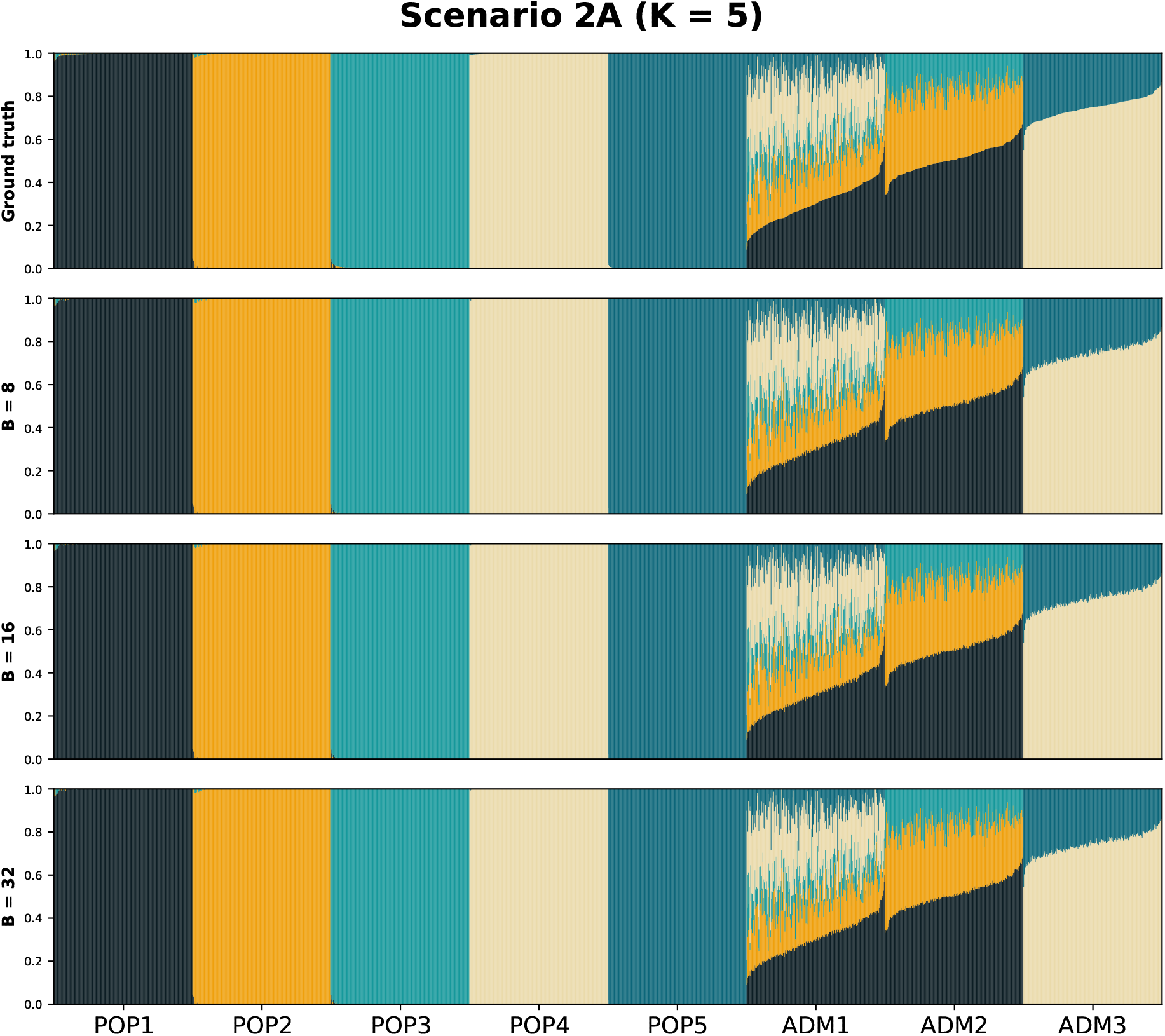
Ancestry plots for the simulated *Scenario 2A* of 2,000 samples for *K* = 5. The top ancestry plot displays the ground truth ancestry proportions from simulated ancestry tracts, followed by the estimates of three different window sizes in `hapla`, *B* = {8, 16, 32}, with *δ* = 0.005.

**Supplementary Figure 5:**
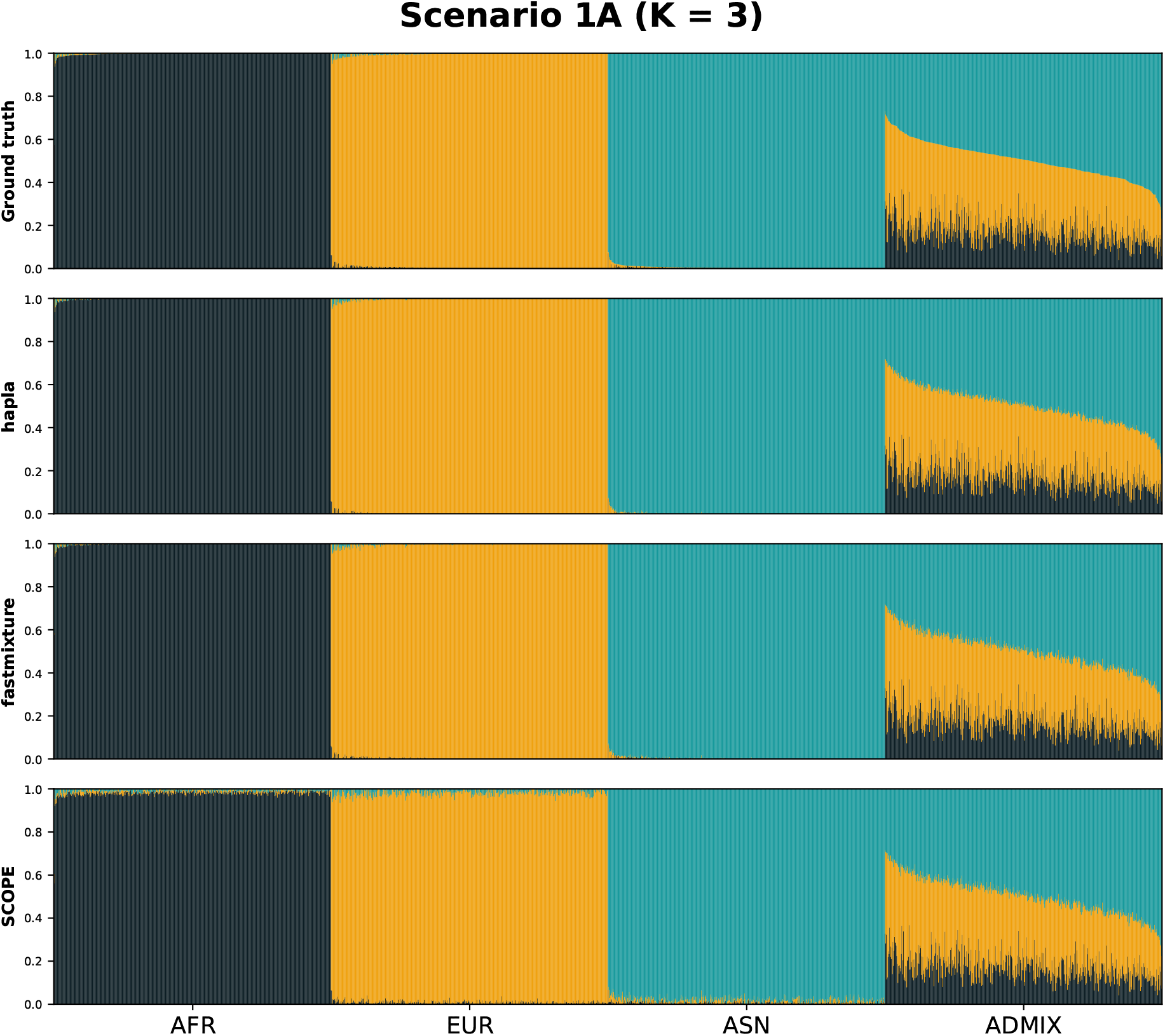
Ancestry plots for the simulated *Scenario 1A* of 2,000 samples for *K* = 5. The top ancestry plot displays the ground truth ancestry proportions from simulated ancestry tracts, followed by the estimates of the three evaluated methods.

**Supplementary Figure 6:**
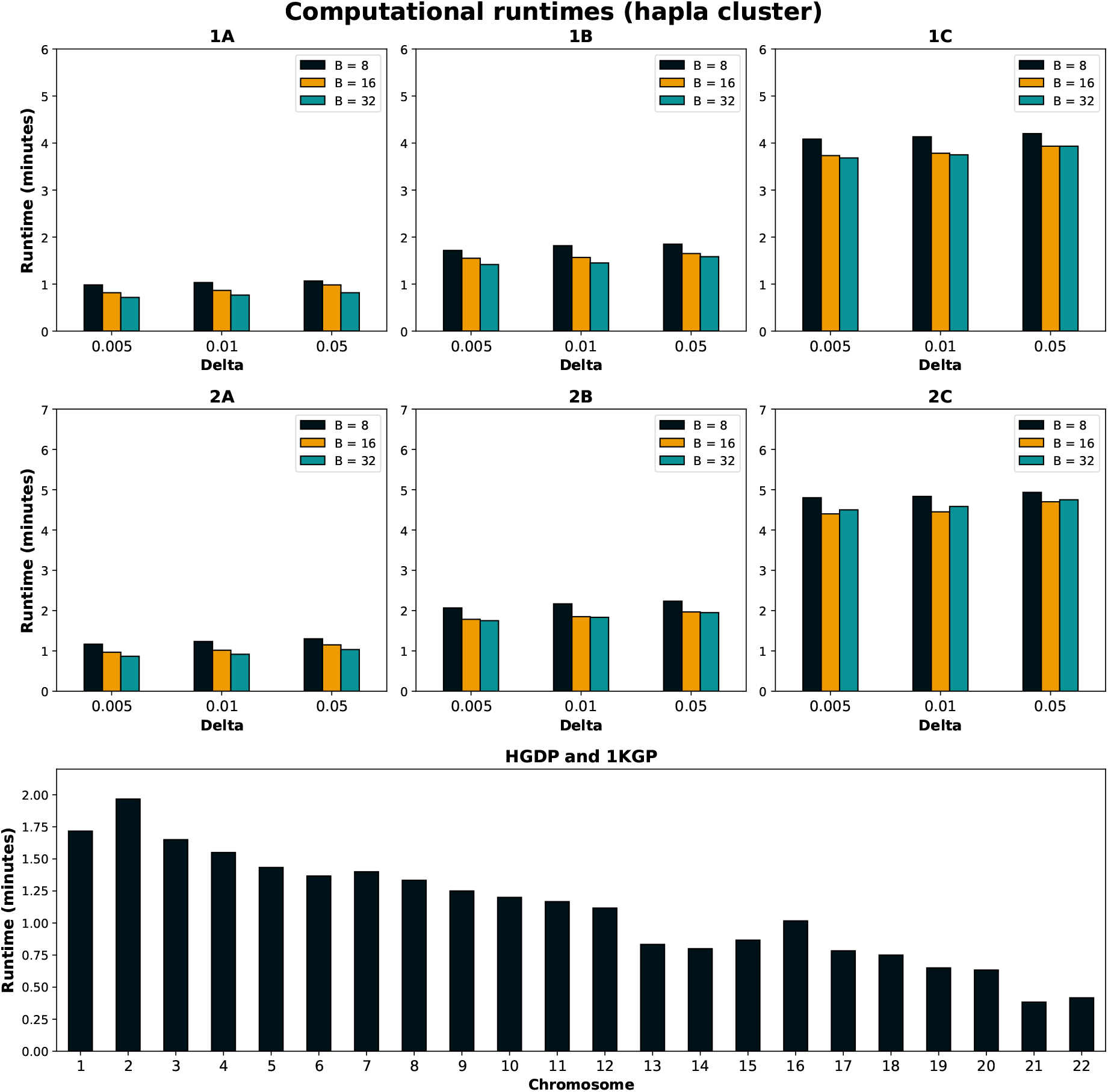
The computational runtime *in minutes* for `hapla cluster` using 44 threads, shown for all parameter combinations in the simulation scenarios and for the harmonized HGDP and 1KGP dataset (only for *B* = 32 and *δ* = 0.005) across all chromosomes. Runtimes include parsing/reading time for the binary VCF (BCF) files using the `cyvcf2` Python library [35]. The simulation scenarios A, B, and C consist of 2,000, 8,000, and 32,000 samples, respectively, while the harmonized HGDP and 1KGP dataset includes 3,406 individuals. The source data for the figure can be found in the Supplemental Data file.

**Supplementary Figure 7:**
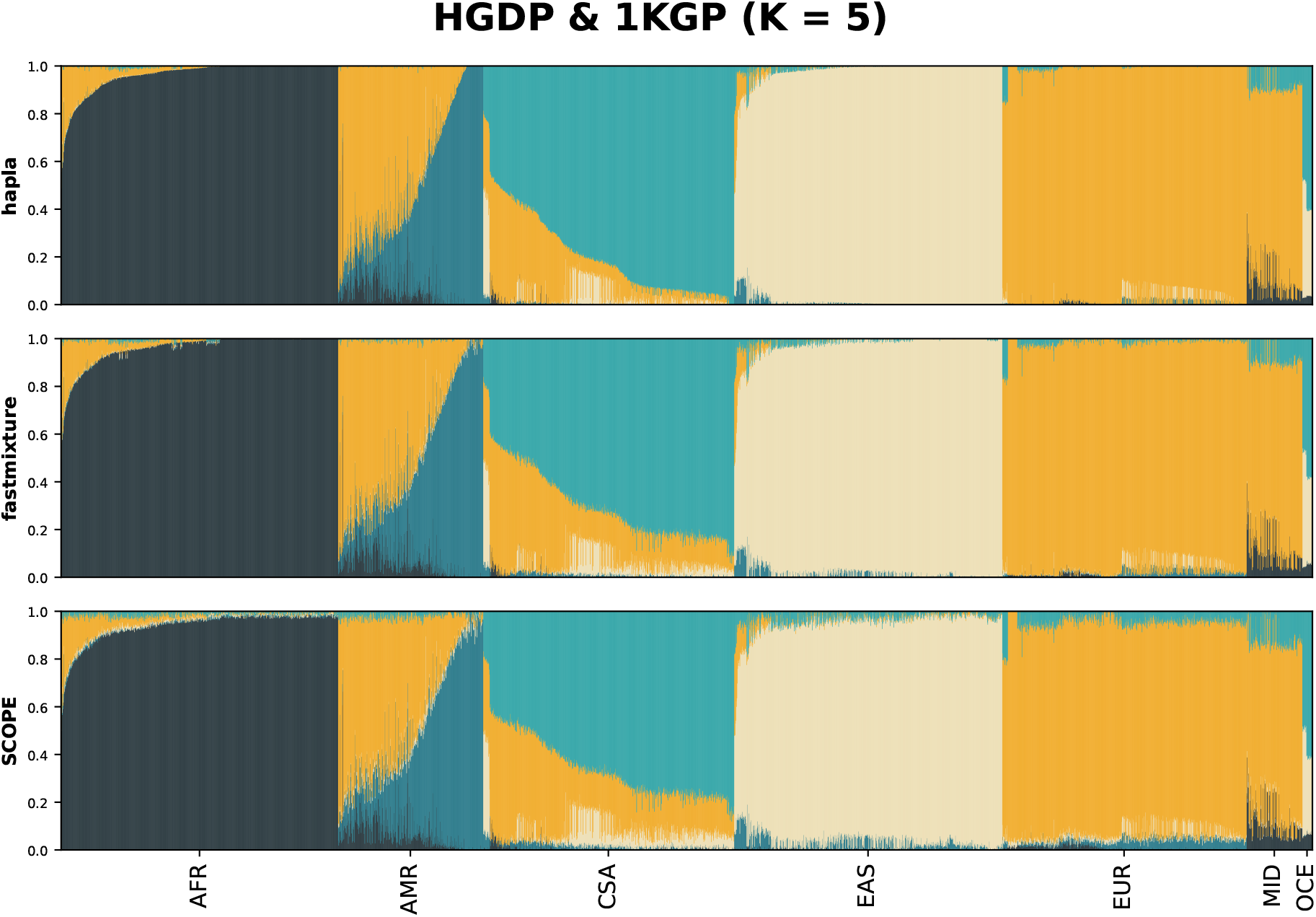
Ancestry plots of 3,406 individuals in the harmonized HGDP and 1KGP dataset for *K* = 5, shown for the three evaluated methods. The order of the individuals is based on metadata labels and the ancestry estimates of `hapla` for *K* = 7.

**Supplementary Figure 8:**
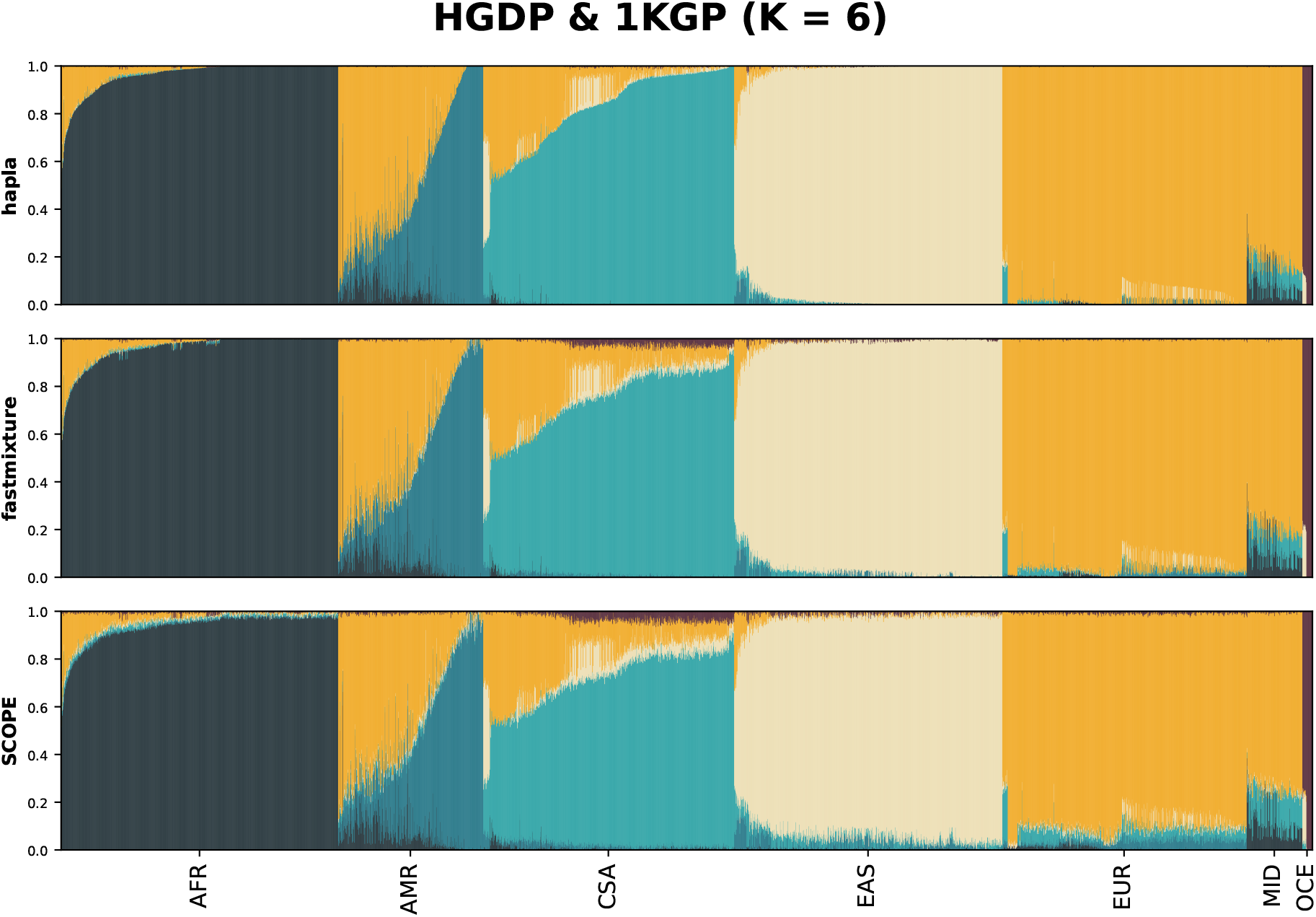
Ancestry plots of 3,406 individuals in the harmonized HGDP and 1KGP dataset for *K* = 6, shown for the three evaluated methods. The order of the individuals is based on metadata labels and the ancestry estimates of `hapla` for *K* = 7.

## Supplementary Tables

### Simulation study

**Supplementary Table 1:** Root mean square error (RMSE) between the ground truth ancestry proportions and the estimated ancestry proportions in the simulated datasets for the different parameter combinations in hapla. Lower values indicate better performance. We have reported the mean RMSE value across five runs, and the standard deviations have been omitted as they are all *<* 1.0 × 10^−5^. The best-performing method in each scenario is highlighted in bold. The source data for the table can be found in the Supplemental Data file.

| Parameters | Scenario 1 |  |  | Scenario 2 |  |  |
| --- | --- | --- | --- | --- | --- | --- |
|  | A | B | C | A | B | C |
| $B = 8, \delta = 0.005$ | 0.00491 | 0.00484 | 0.00502 | 0.00383 | 0.00387 | 0.00390 |
| $B = 16, \delta = 0.005$ | **0.00473** | **0.00453** | 0.00469 | 0.00358 | 0.00349 | 0.00349 |
| $B = 32, \delta = 0.005$ | 0.00490 | 0.00459 | **0.00464** | **0.00354** | **0.00343** | **0.00340** |
| $B = 8, \delta = 0.01$ | 0.00494 | 0.00487 | 0.00506 | 0.00392 | 0.00396 | 0.00399 |
| $B = 16, \delta = 0.01$ | 0.00477 | 0.00457 | 0.00474 | 0.00366 | 0.00358 | 0.00359 |
| $B = 32, \delta = 0.01$ | 0.00493 | 0.00463 | 0.00470 | 0.00366 | 0.00354 | 0.00352 |
| $B = 8, \delta = 0.05$ | 0.00521 | 0.00515 | 0.00533 | 0.00431 | 0.00447 | 0.00447 |
| $B = 16, \delta = 0.05$ | 0.00508 | 0.00498 | 0.00517 | 0.00427 | 0.00435 | 0.00434 |
| $B = 32, \delta = 0.05$ | 0.00536 | 0.00516 | 0.00532 | 0.00459 | 0.00457 | 0.00456 |

**Supplementary Table 2:** Root mean square error (RMSE) between the ground truth ancestry proportions and the estimated ancestry proportions of ***the admixed populations only*** in the simulated datasets. Lower values indicate better performance. We have reported the mean RMSE value across five runs, and the standard deviations have been omitted as they are all *<* 1.0 × 10^−5^. The best-performing method in each scenario is highlighted in bold. `hapla` is based on *B* = 32 and *δ* = 0.005. The source data for the table can be found in the Supplemental Data file.

| Method | Scenario 1 |  |  | Scenario 2 |  |  |
| --- | --- | --- | --- | --- | --- | --- |
|  | A | B | C | A | B | C |
| SCOPE | 0.01408 | 0.01448 | 0.01423 | 0.01153 | 0.01196 | 0.01189 |
| fastmixture | 0.01221 | 0.01232 | 0.01227 | 0.00841 | 0.00864 | 0.00870 |
| hapla | **0.00821** | **0.00801** | **0.00802** | **0.00545** | **0.00539** | **0.00532** |

### Harmonized HGDP and 1KGP data

**Supplementary Table 3:** Mean log-likelihood across five runs in the harmonized HGDP and 1KGP dataset for three different values of *K*, with standard deviations reported in parentheses. Note that log-likelihoods of `hapla` are not comparable with the two other SNP-based approaches. `hapla` was run with *B* = 32 and *δ* = 0.005. The source data for the table can be found in the Supplemental Data file.

| Method | Log-likelihoods |  |  |
| --- | --- | --- | --- |
| | $K = 5$ | $K = 6$ | $K = 7$ |
| SCOPE | -17826107180.6 (4745.1) | -17791745046.5 (6424.8) | -17783061309.1 (1202.3) |
| fastmixture | -17820987921.4 (1.2) | -17786024270.6 (2.1) | -17764854092.0 (4.2) |
| hapla | -1985267046.6 (1.9) | -1979789715.7 (6.0) | -1975446168.8 (29.6) |

**Supplementary Table 4:** Akaike information criterion (AIC) in the harmonized HGDP and 1KGP dataset for three different values of *K*. Lower values indicate a more appropriate model. `hapla` was run with *B* = 32 and *δ* = 0.005. The source data for the table can be found in the Supplemental Data file.

| Method | AIC |  |  |
| --- | --- | --- | --- |
| | $K = 5$ | $K = 6$ | $K = 7$ |
| SCOPE | 35710945029 | 35653968257 | **35648348278** |
| fastmixture | 35700706511 | 35642526705 | **35611933844** |
| hapla | 3993325831 | 3986930879 | **3982803496** |

### Computational runtimes (ancestry estimation)

**Supplementary Table 5:** Mean computational runtime *in minutes* across five runs using 44 threads in the simulated datasets for different parameter combinations in hapla, with standard deviations shown in parentheses. Lower values indicate faster runtimes. The source data for the table can be found in the Supplemental Data file.

| Parameters | Scenario 1 |  |  | Scenario 2 |  |  |
| --- | --- | --- | --- | --- | --- | --- |
|  | A | B | C | A | B | C |
| $B = 8, \delta = 0.005$ | 0.21 (0.03) | 0.68 (0.05) | 2.58 (0.15) | 0.25 (0.03) | 0.89 (0.09) | 3.36 (0.61) |
| $B = 16, \delta = 0.005$ | 0.12 (0.01) | 0.39 (0.06) | 1.57 (0.08) | 0.15 (0.01) | 0.49 (0.03) | 1.94 (0.13) |
| $B = 32, \delta = 0.005$ | **0.07 (0.01)** | 0.25 (0.02) | 0.98 (0.14) | 0.10 (0.02) | 0.28 (0.01) | 1.16 (0.09) |
| $B = 8, \delta = 0.01$ | 0.19 (0.01) | 0.65 (0.05) | 2.65 (0.16) | 0.23 (0.02) | 0.83 (0.09) | 3.28 (0.36) |
| $B = 16, \delta = 0.01$ | 0.11 (0.01) | 0.38 (0.02) | 1.44 (0.11) | 0.15 (0.01) | 0.50 (0.05) | 1.81 (0.13) |
| $B = 32, \delta = 0.01$ | **0.07 (0.01)** | 0.25 (0.03) | **0.88 (0.06)** | 0.08 (0.01) | 0.28 (0.02) | 1.10 (0.09) |
| $B = 8, \delta = 0.05$ | 0.20 (0.04) | 0.66 (0.07) | 2.65 (0.15) | 0.26 (0.02) | 0.85 (0.06) | 3.58 (0.56) |
| $B = 16, \delta = 0.05$ | 0.11 (0.01) | 0.38 (0.05) | 1.48 (0.14) | 0.15 (0.01) | 0.50 (0.03) | 1.75 (0.17) |
| $B = 32, \delta = 0.05$ | **0.07 (0.01)** | **0.23 (0.02)** | 0.91 (0.10) | **0.07 (0.01)** | **0.27 (0.03)** | **1.08 (0.09)** |

**Supplementary Table 6:** Mean computational runtime *in minutes* across five runs using 44 threads in the simulated datasets, with standard deviations reported in parentheses. Lower values indicate faster runtimes. `hapla` is based on *B* = 32 and *δ* = 0.005. The source data for the table can be found in the Supplemental Data file.

| Method | Scenario 1 |  |  | Scenario 2 |  |  |
| --- | --- | --- | --- | --- | --- | --- |
|  | A | B | C | A | B | C |
| SCOPE | 0.15 (0.00) | 0.39 (0.00) | 1.25 (0.01) | 0.29 (0.04) | 0.66 (0.07) | 2.45 (0.08) |
| fastmixture | 0.77 (0.05) | 3.21 (0.20) | 13.06 (1.68) | 1.06 (0.17) | 3.52 (0.43) | 17.70 (1.65) |
| **hapla** | **0.07 (0.01)** | **0.25 (0.02)** | **0.98 (0.14)** | **0.10 (0.02)** | **0.28 (0.01)** | **1.16 (0.09)** |

**Supplementary Table 7:** Mean computational runtime *in minutes* across five runs using 44 threads in the harmonized HGDP and 1KGP dataset for three different *K*, with standard deviations reported in parentheses. Lower values indicate faster runtimes. `hapla` is based on *B* = 32 and *δ* = 0.005. The source data for the table can be found in the Supplemental Data file.

| Method | HGDP and 1KGP |  |  |
| --- | --- | --- | --- |
| | $K = 5$ | $K = 6$ | $K = 7$ |
| SCOPE | 12.46 (2.01) | 14.65 (1.17) | 30.29 (5.12) |
| fastmixture | 48.25 (2.66) | 51.97 (2.61) | 116.20 (3.63) |
| **hapla** | **2.34 (0.11)** | **2.62 (0.15)** | **6.20 (0.40)** |

